# Maftivimab monotherapy is effective in a non-human primate model of Bundibugyo virus disease

**DOI:** 10.64898/2026.10.13.758840

**Authors:** Meike Dittmann, Dylan M. Johnson, Micah Rapp, Alida Coppi, Flonza Isa, Dharani Ajithdoss, Anbo Zhou, Benjamin O. Fulton, Ranu Basu, Anna K. Kania, Kristen E. Pascal, Robert LeDesma, George Ehrlich, Ashique Rafique, Victoria Marzec, Jiacheng Yuan, Ruiqi Liu, Kendra Alfson, Tessa Moir-Savitz, Stanton Gray, Greg Ippolito, Cory Hallam, Jean Patterson, Alina Baum, Veronica Mas Casullo, Richard Copin, Viraj Kulkarni, Matthew C. Franklin, Christos A. Kyratsous, Jamie M. Orengo

## Abstract

The 2026 outbreak of Bundibugyo ebolavirus (BDBV) in the Democratic Republic of the Congo and Uganda has created an urgent need for treatments for BDBV disease, yet the only FDA-approved ebolavirus therapeutics, the monoclonal antibody (mAb) cocktail Inmazeb® and the mAb Ebanga®, are indicated for Zaire ebolavirus (EBOV). Maftivimab (REGN3479), the most potent neutralizing component of Inmazeb®, targets ebolavirus species GPs broadly; however, how it engages BDBV GP, whether it neutralizes BDBV-2026, and whether it protects against BDBV disease *in vivo* are unknown. Here we show that Maftivimab binds to BDBV-2007 GP with high affinity and potently neutralizes pseudoviruses carrying GP from a 2007 reference strain and the 2026 outbreak lineage. Additionally, a cryo-electron microscopy structure of Maftivimab Fab bound to trimeric BDBV-2007 GP superimposes closely on the EBOV GP–Maftivimab complex; importantly, all GP contact residues are conserved between the 2007 and 2026 BDBV sequences. Finally, in cynomolgus macaques challenged with BDBV-2007, a single dose of Maftivimab monotherapy given 7 days after infection, after the onset of clinical disease, rescued 6 of 6 animals, whereas 5 of 6 placebo-treated animals succumbed. These findings demonstrate that Maftivimab neutralizes BDBV, binds to its epitope on BDBV GP and is efficacious in treating established BDBV disease in the macaque model. The data support its evaluation in clinical trials during the ongoing or future outbreak.

## Introduction

Since its emergence in the Democratic Republic of the Congo and Uganda in 2026, the current outbreak of *Orthoebolavirus bundibugyoense* (BDBV) has grown into the largest recorded BDBV outbreak and second largest filovirus outbreak to date. Currently, there are no therapeutics or vaccines approved for the treatment or prevention of BDBV^1^. Projections suggest this outbreak could eclipse the most severe filovirus outbreak^2^, the 2013-2016 West African *Orthoebolavirus zairense* (EBOV) epidemic. In the absence of approved countermeasures, there is an urgent public health need to identify potential BDBV treatments and interrogate their efficacy *in vitro* and in non-human primate (NHP) challenge models, to support further evaluation in controlled clinical trials.

Monoclonal antibodies (mAbs) specific for EBOV glycoprotein (GP) are the only FDA-approved treatments for ebolavirus disease; however, they are indicated only for EBOV and not for any other ebolavirus species^3,4^. We previously showed that Inmazeb®, a cocktail of three human mAbs (Atoltivimab, Odesivimab, and Maftivimab), was efficacious against EBOV in NHPs and then in humans in the Pamoja Tulinde Maisha (PALM) clinical trial, conducted during the 2018–2020 EBOV outbreak^5,6^. This led to Inmazeb® becoming the first FDA-approved treatment for EBOV infection and to its inclusion on the WHO List of Prequalified Products^3,7^. A second mAb therapeutic, mAb114 (Ebanga®), was also evaluated in the PALM clinical trial and is likewise approved for the treatment of EBOV disease^4^. The extent to which these GP-directed therapeutics retain efficacy against the more distantly related BDBV — whose GP shares approximately 65% overall amino acid identity with EBOV GP, driven largely by divergence in the hypervariable mucin-like domain^8^ — remains an open and clinically urgent question in the context of the ongoing 2026 BDBV outbreak.

Maftivimab (REGN3479), a human IgG1κ mAb generated using the VelocImmune® technology^9,10^, is the broadest and most potent neutralizing component of Inmazeb®^6^. Cryo-EM structures of the Inmazeb®–EBOV GP complex show that Maftivimab binds a quaternary epitope of the trimeric pre-fusion GP complex spanning the GP1 internal fusion loop and GP2 heptad repeat 1 (HR1) across adjacent GP monomers and prevents entry of the virus into susceptible cells^11^. However, it is unknown whether Maftivimab binds to the equivalent epitope on the BDBV GP, whether that epitope is conserved in the 2026 outbreak lineage, and whether Maftivimab can neutralize BDBV-2026 and protect against BDBV disease *in vivo*. Here, we report quantitative binding kinetics and pseudovirus neutralization data for Maftivimab against BDBV, the cryo-EM structure of Maftivimab Fab in complex with trimeric BDBV GP, and *in vivo* efficacy in the cynomolgus macaque NHP challenge model, to demonstrate that Maftivimab is effective against BDBV in preclinical models.

## Results

### Maftivimab binds BDBV GP with low-nanomolar affinity and neutralizes pseudovirus carrying GP from 2007 and 2026 BDBV lineages

To evaluate and quantify Maftivimab binding to BDBV-2007 GP, we measured binding kinetics by surface plasmon resonance (Biacore). Maftivimab bound to BDBV-2007 GP trimer with an equilibrium dissociation constant (K_D)_ of 7.88E-09 M and a dissociation half-life of t½ = 96.9 min at 37 °C (**Figure 1a**). By comparison, Maftivimab bound EBOV GP trimer with a KD of 1.98E-09 M and a t½ of 172.3 min, consistent with previously reported values^11^. These data demonstrate that Maftivimab binds BDBV GP with low-nanomolar affinity and a dissociation half-life of more than 1.5 hours, similar to its binding kinetics to EBOV GP.

**Figure 1.**
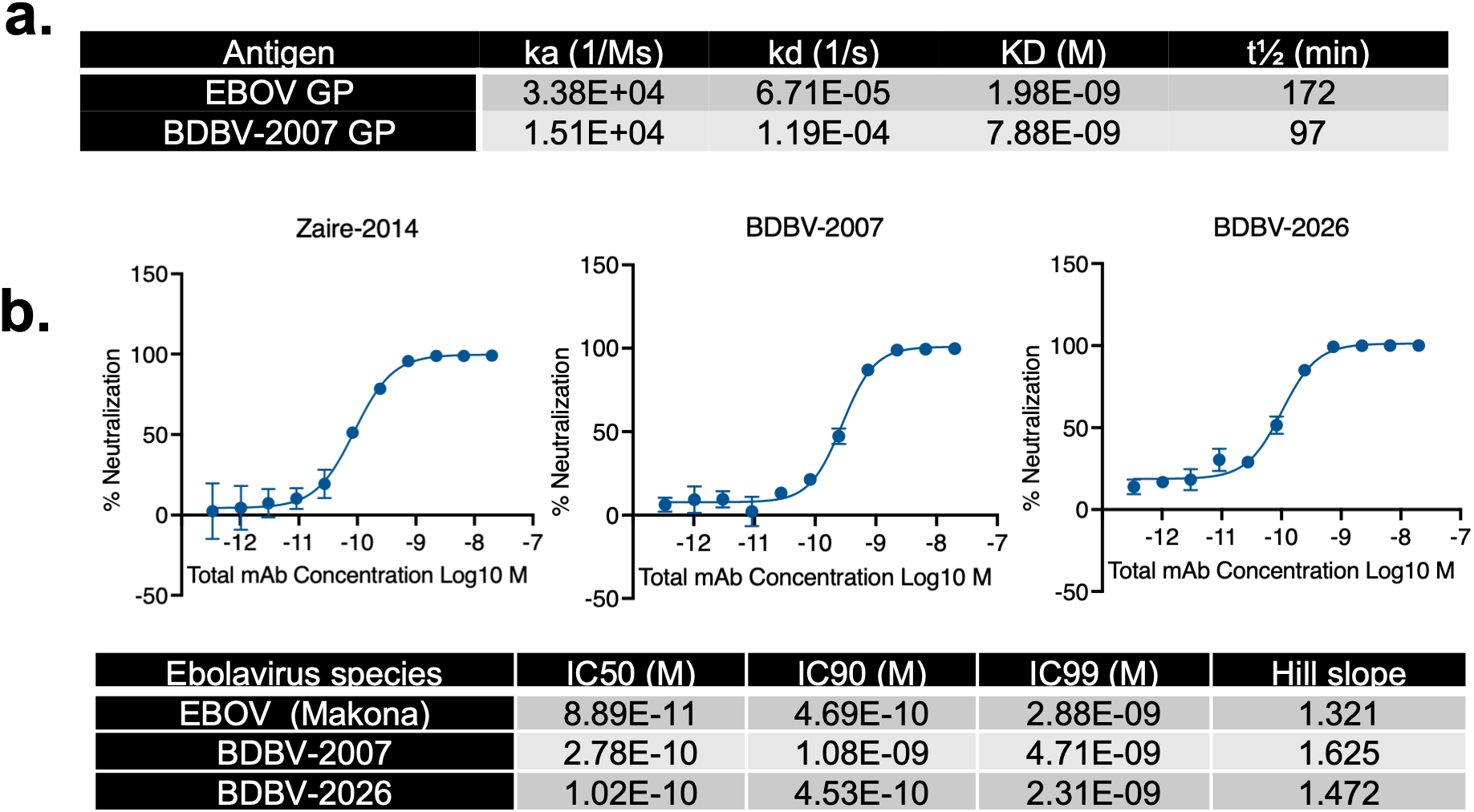
Binding kinetics and neutralization potency of Maftivimab (REGN3479) against BDBV. (a) Surface plasmon resonance binding kinetics of Maftivimab against EBOV (GenBank AHX24649.2) and BDBV-2007 (GenBank YP_003815435.1) GP trimers (equilibrium dissociation constant, KD; dissociation half-life, t½) at 37 °C. (b) Pseudovirus neutralization curves and potency (IC50, IC90, IC99, Hill slope) of Maftivimab against VSV pseudovirus bearing full-length GP from EBOV (GenBank AHX24649.1), BDBV-2007 (GenBank YP_003815435.1), and BDBV-2026 (PP_006XCJJ.1; published May 17, 2026). VSV, replication-incompetent vesicular stomatitis virus encoding a luciferase reporter; mAb, monoclonal antibody; SEM, standard error of the mean. An 11-point mAb dilution series was incubated with pseudovirus in triplicate and applied to Vero cells for 24 h. Infection was quantified by luciferase activity, and % neutralization was calculated relative to uninfected and infected no-mAb control wells. Points represent mean values; error bars indicate SEM.

We have previously shown that Maftivimab neutralizes authentic EBOV and BDBV-2007^11^. To determine whether Maftivimab retains neutralizing activity against the 2026 BDBV lineage, for which an authentic isolate is not yet available, we tested the ability of Maftivimab to neutralize replication-incompetent vesicular stomatitis virus (VSV) pseudovirus carrying GP from EBOV, BDBV-2007 or BDBV-2026. As previously demonstrated^11^, Maftivimab potently neutralized EBOV GP pseudovirus (IC50 = 8.89E-11 M). Maftivimab also neutralized pseudoviruses carrying GP from either BDBV with high, sub-nanomolar potency (IC50 = 2.78E-10 M and 1.02E-10 M for 2007 and 2026, respectively; **Figure 1b**), demonstrating that it is a broad neutralizer with activity extending to BDBV. Neutralization potency was similar between BDBV-2007 and BDBV-2026. Together, these data show that Maftivimab binds BDBV GP with high affinity and potently neutralizes both the 2007 and 2026 lineages. We next determined how Maftivimab engages BDBV GP.

### Cryo-EM structure of Maftivimab Fab in complex with trimeric BDBV-2007 GP reveals canonical quaternary epitope engagement

To determine whether Maftivimab engages the EBOV-homologous epitope on the BDBV glycoprotein, we determined the cryo-EM structure of Maftivimab Fab in complex with trimeric BDBV-2007 GP (YP_003815435.1) at 2.9 Å resolution. Maftivimab engages a quaternary BDBV GP epitope that bridges the internal fusion loop of one GP monomer to a GP1/GP2-containing site in an adjacent monomer (**Figure 2a**).

**Figure 2.**
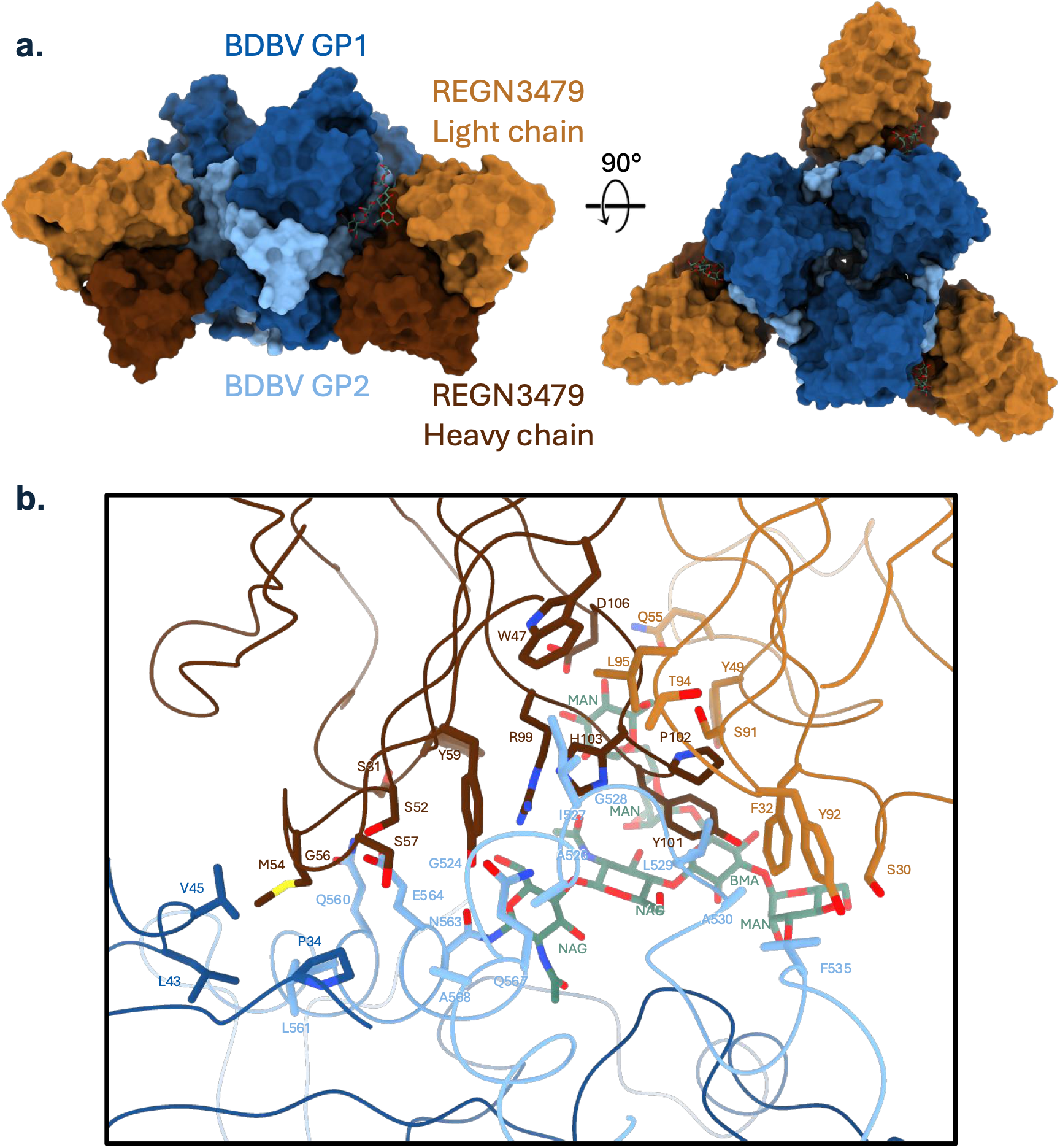
Cryo-EM structure of Maftivimab (REGN3479) Fab bound to trimeric BDBV-2007 glycoprotein. (a) Surface representation of the Fab–GP complex in a front view and a 90°-rotated top-down view, showing three copies of the REGN3479 Fab (light chain, orange; heavy chain, brown) engaging the BDBV-2007 glycoprotein trimer (GP1, dark blue; GP2, light blue) at the internal fusion loop. (b) Close-up of the Fab–GP interface showing REGN3479 paratope and BDBV-2007 GP epitope residues, defined as residues with at least 15 Å² of buried surface area. The glycan at N563 is also shown.

Direct contact residues on GP1 include P34, L43, and V45, while direct contacts on GP2 span G524, A526, I527–A530 (IGLA motif), F535, and HR1 residues Q560, L561, E564, Q567, and A568, as well as Manα1-6Manα1-6Manβ1-4GlcNAcβ1-4GlcNAc in the glycan at N563 (**Figure 2b**). Paratope residues from all six antibody complementarity-determining regions (CDRs) are involved in contacting GP; CDRH1 interacts with some HR1 residues as well as the terminal Manα1-6 of the N563 glycan, CDRH2 makes extensive contacts with HR1, GP1, and flanks one side of the fusion loop, and CDRH3 interacts with the fusion loop and almost all saccharides in the N563 glycan. On the light chain, CDRL1 and L3 also flank the fusion loop, whereas CDRL2 interacts with two Mannose moieties on the N563 glycan (**Figure 2b**). Together, these data show that Maftivimab engages BDBV-2007 GP through a quaternary epitope that spans the internal fusion loop, HR1 and the N563 glycan, with all six CDRs contributing.

### Maftivimab engages a similar epitope on BDBV GP and EBOV GP

Having defined how Maftivimab engages BDBV-2007 GP, we next asked whether this binding mode matches that on EBOV GP. We thus superimposed our BDBV-2007 GP:Maftivimab structure onto the Maftivimab:GP portion of the previously reported EBOV GP:Inmazeb® structure^11^. The two complexes align closely (C-alpha Root Mean Square Deviation = 1.3 Å), and Maftivimab binds a similar quaternary epitope, contacting corresponding residue positions, on both glycoproteins (**Figure 3a, b**).

**Figure 3.**
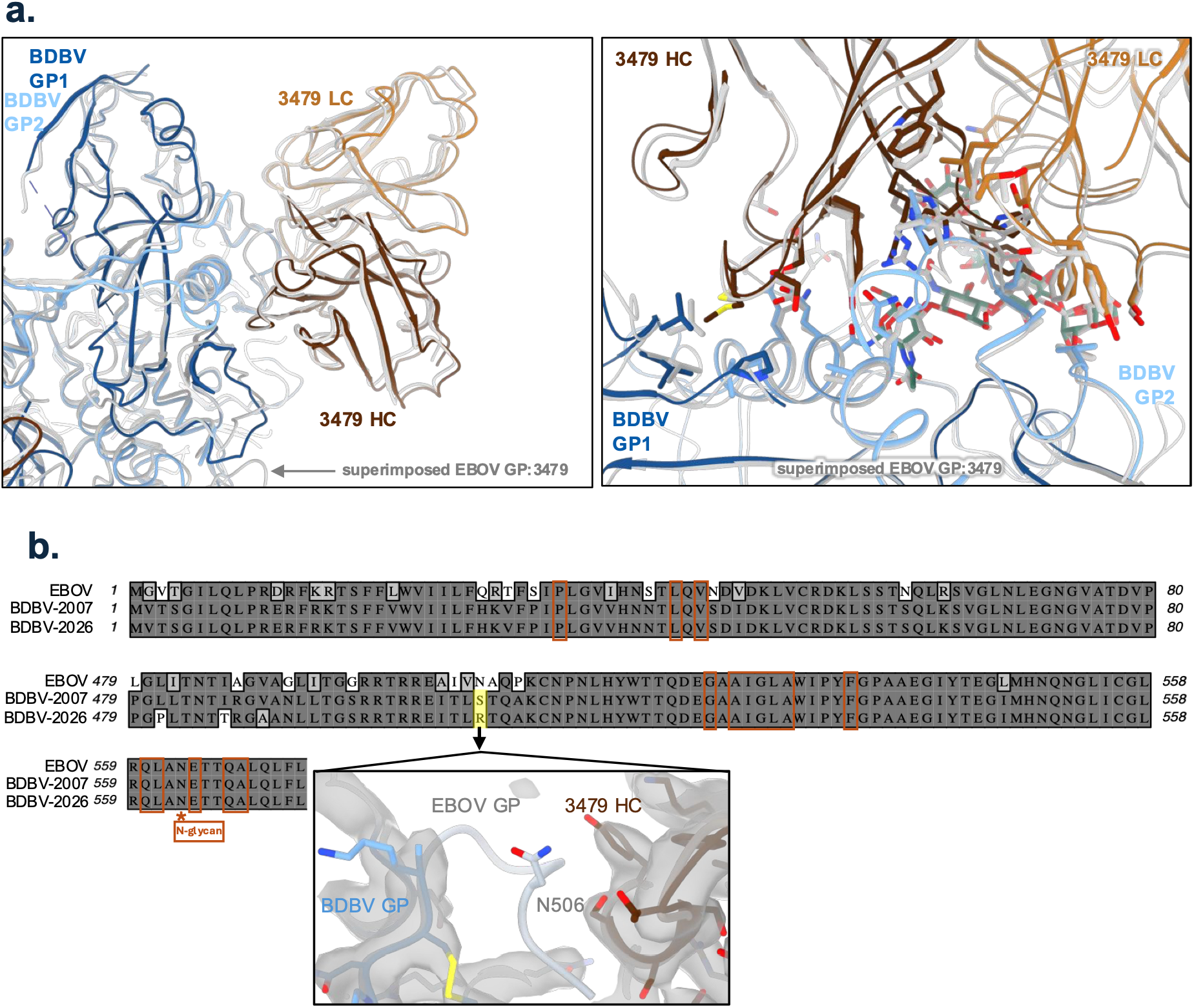
Maftivimab (REGN3479) engages equivalent epitopes on BDBV GP and EBOV GP. (a) Superposition of the BDBV-2007 GP–REGN3479 structure and the previously reported EBOV GP–REGN3479 structure (PDB 7TN9)^11^. In the BDBV-2007 GP–REGN3479 structure, the REGN3479 Fab light chain is orange, the heavy chain is brown, and the BDBV-2007 GP trimer is dark blue (GP1) and light blue (GP2). The EBOV GP–REGN3479 structure is shown entirely in gray. Left, overview of the Fab–GP interface; right, close-up of the epitope. Root mean square deviation (RMSD) = 1.3 Å. (b) Sequence alignment of EBOV (Makona isolate Kissidougou-C15, GenBank AHX24649.2), BDBV-2007 (isolate Butalya-811250, GenBank YP_003815435.1) and BDBV-2026 (Uganda index case, Pathoplexus PP_006XCJJ.1)^12^ GP; for conciseness, only the sequence regions containing REGN3479 contact residues are depicted. Brown boxes denote REGN3479 contact residues in our BDBV-2007 complex structure; the star marks residue N563, which is not itself a REGN3479 contact but carries an N-linked glycan that is directly contacted. Inset, map and model overlay of the BDBV-2007 and EBOV structures focused on residue N506; density for this residue is not present in the BDBV-2007 GP–REGN3479 reconstruction.

To assess whether the contacts seen in our BDBV-2007 structure are retained in BDBV-2026, we aligned EBOV (Makona), BDBV-2007 (Butalya-811250) and BDBV-2026 (Uganda index case)^12^ GP sequences across the regions targeted by Maftivimab (**Figure 3b**). All direct contact residues observed in our BDBV-2007 complex structure are 100% conserved between EBOV, BDBV-2007 and BDBV-2026 GP. Residues 504-509 are not conserved among the three glycoproteins (IVNAQP in EBOV, TLSTQA in BDBV-2007, TLRTQA in BDBV-2026) and residue 506 in this region differs between BDBV-2007 and BDBV-2026 (**Figure 3b**). While this region of GP makes weak contacts with Maftivimab in the EBOV GP:Inmazeb® complex^11^, we do not observe any EM density for these amino acids in our BDBV-2007 GP:Maftivimab complex, although the surrounding GP and Maftivimab residues are generally well defined (**Figure 3b, inset**). This lack of EM density indicates that there are no interactions between BDBV-2007 GP residues 504-509 and Maftivimab. In the published EBOV GP:Inmazeb® structure, N506 lies close to the Maftivimab heavy chain but makes no direct contact (**Figure 3b, inset**), implying that this position is not critical for binding and neutralization activity.

### The Maftivimab epitope is conserved across 2026 BDBV outbreak sequences

To determine whether the Maftivimab epitope is conserved among BDBV sequences from the 2026 outbreak, we analyzed per-position GP conservation across 60 available GenBank sequences (**Figure 4**). All 15 Maftivimab contact residues and the 25 neighboring residues within 5 Å were invariant, indicating a fully conserved epitope. Together with the similar neutralization potency of BDBV-2007 and BDBV-2026, this indicates that Maftivimab is expected to remain active against viruses circulating in the current outbreak.

**Figure 4.**
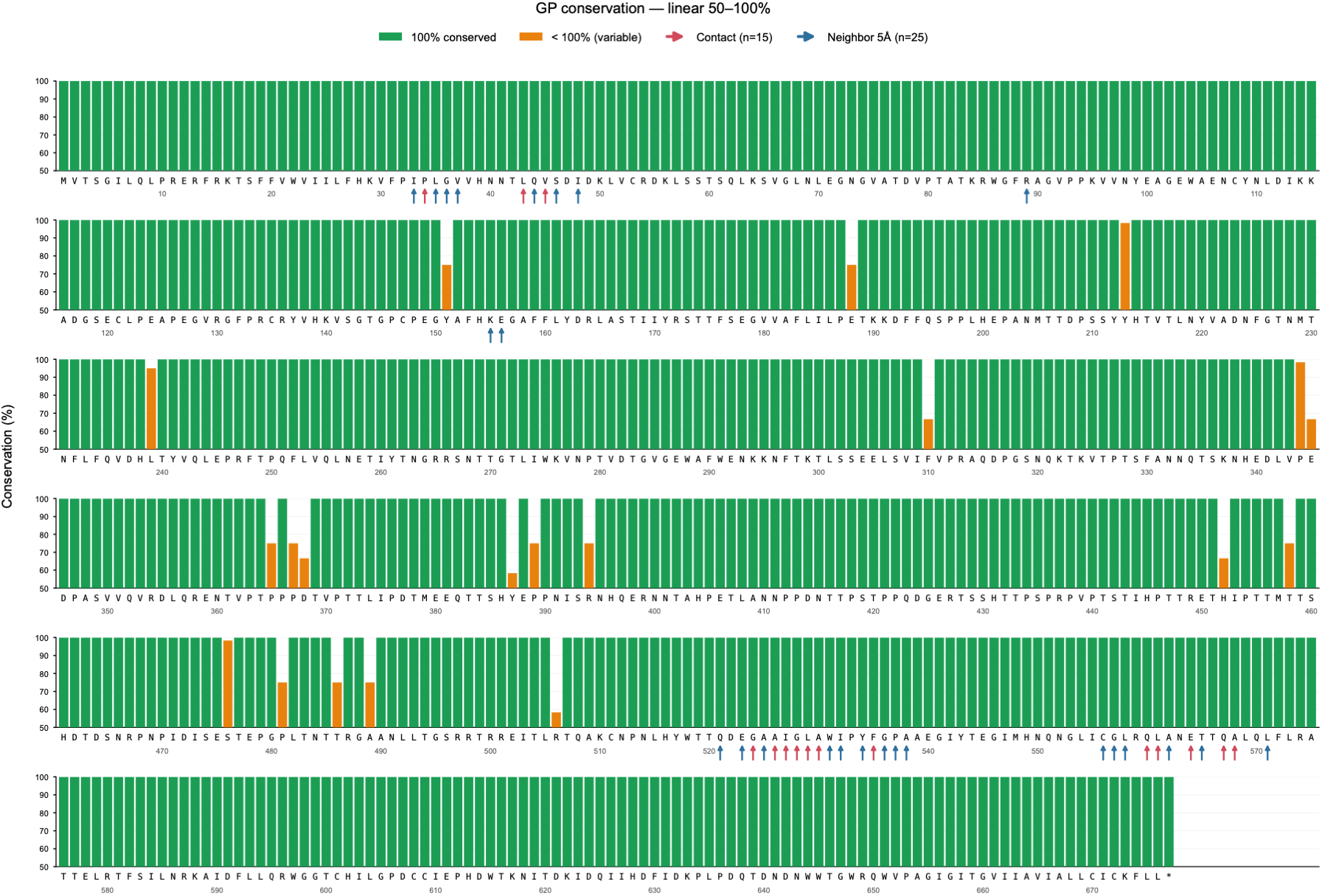
Conservation of Maftivimab binding residues within 2026 BDBV outbreak sequences. Per-position conservation of BDBV GP across 2026 outbreak sequences was calculated from 60 high-quality glycoprotein sequences retrieved from NCBI GenBank, y axis truncated at 90–100%. Green bars, 100% conserved positions; orange bars, variable positions (<100%). Red arrows, Maftivimab contact residues (n = 15); blue arrows, neighboring residues within 5 Å (n = 25).

### Maftivimab monotherapy initiated after onset of clinical disease protects cynomolgus macaques from lethal BDBV challenge

To determine whether the *in vitro* activity of Maftivimab translates to protection *in vivo*, we evaluated Maftivimab monotherapy in a lethal BDBV challenge model in cynomolgus macaques (*Macaca fascicularis*) using BDBV-2007. NHPs were challenged intramuscularly with 1000 plaque-forming units (PFUs) of BDBV-2007, stratified into treatment groups by sex and viral load measured by RT-qPCR at 4 days post infection (DPI) (**Figure 5a**), and treated intravenously with saline (placebo) or 150 mg/kg Maftivimab (n = 6 per group), matching the total antibody dose of the Inmazeb® cocktail (50 mg/kg each of Atoltivimab, Odesivimab and Maftivimab), at 7 DPI. A significant survival benefit was observed with all six Maftivimab-treated animals (100%) compared with one of six placebo-treated animals (16.7% survival) reaching the planned end of the study at 28 DPI (**Figure 5b**).

**Figure 5.**
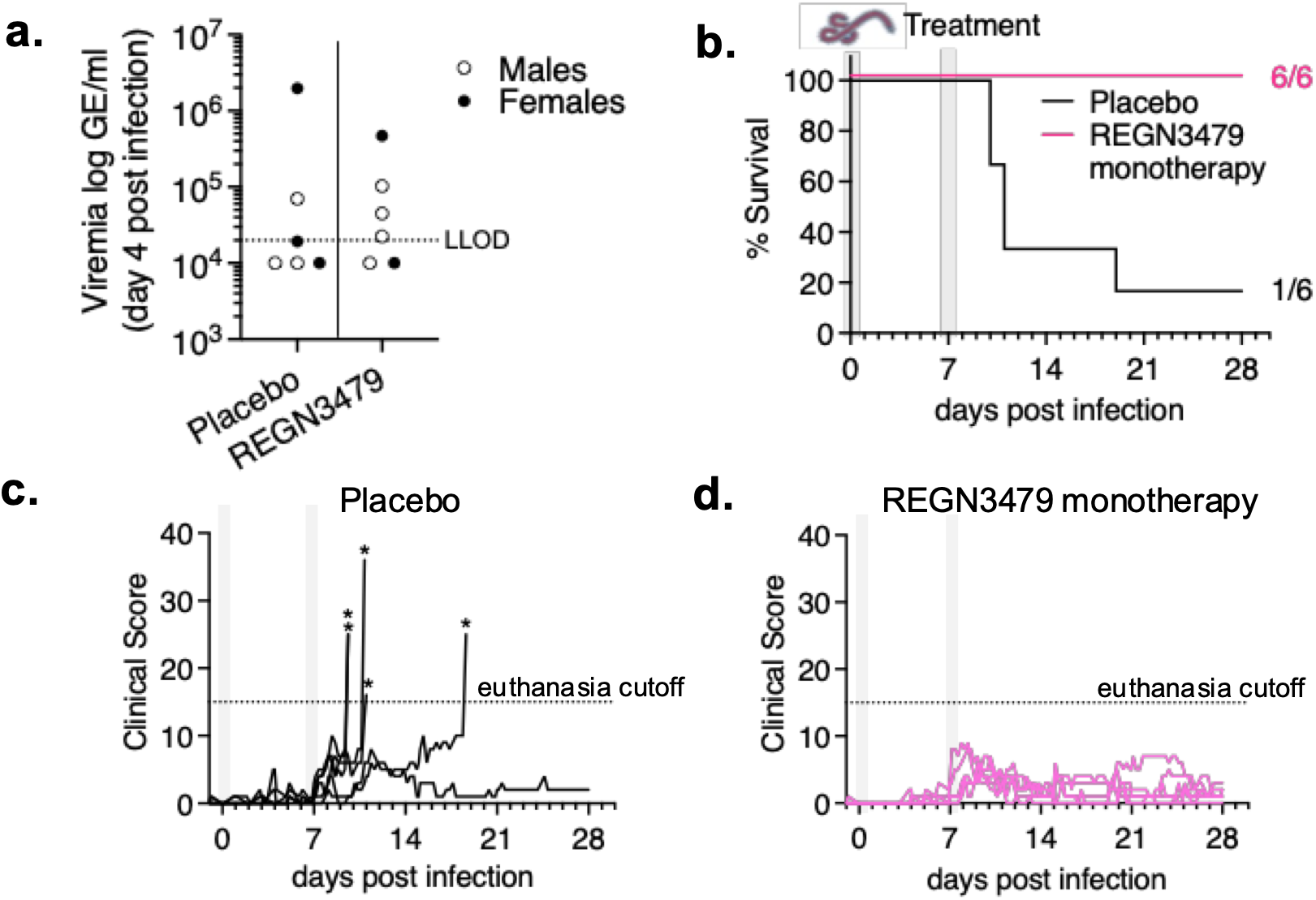
*In vivo* efficacy of Maftivimab (REGN3479) monotherapy in cynomolgus macaques challenged with BDBV. Cynomolgus macaques (nonhuman primates, NHPs) were infected intramuscularly (IM) with 1000 plaque-forming units (PFU) of BDBV-2007 on day 0 (DPI, days post infection), stratified by sex and 4 DPI viral load, and treated intravenously (IV) on 7 DPI with saline (placebo) or 150 mg/kg Maftivimab (n = 6 per group). (a) Viremia (log GE/mL) on 4 DPI, used for stratification. Open symbols, males; closed symbols, females. The dotted line represents the lower limit of detection (LLOD) based on the detection of control plasmid. Samples with both technical replicates below the LLOD are plotted as half the LLOD. Indeterminate samples (with one replicate above the LLOD and one below) are plotted as the average of the replicate values, and those NHPs were considered PCR-positive for stratification purposes. (b) Survival through 28 DPI; survival at study end was compared between groups using a two-sided Boschloo’s exact test (p = 0.0039). Gray bars indicate infection (0 DPI) and treatment (7 DPI). Non-survival was defined as terminal illness based on cumulative clinical scoring. (c, d) Clinical scores of placebo-treated (c) and Maftivimab-treated (d) animals, assessed daily through 28 DPI. Dotted line, euthanasia cutoff. Asterisks, day of euthanasia. GE, genome equivalents.

Five of six placebo-treated NHPs met humane early euthanasia criteria, with clinical scores rising sharply before death (**Figure 5c**). Notably, all NHPs meeting early euthanasia criteria scored for reduced responsiveness (defined by moderate to severe reduced responses to external stimuli and/or assuming of a prostrate position with or without the ability to rise if stimulated). In contrast, clinical scores of Maftivimab-treated NHPs remained below the euthanasia cutoff throughout the study and moderate to severely reduced responsiveness was never observed in this group (**Figure 5d**).

Together, these data show that a single dose of Maftivimab monotherapy, given after the onset of clinical BDBV symptoms, protects macaques from lethal disease. Since Maftivimab neutralizes BDBV-2007 and BDBV-2026 with similar potency (**Figure 1b**), protection against BDBV-2007 is expected to extend to the 2026 lineage.

## Discussion

Our experimental data show that Maftivimab (REGN3479) retains binding to its quaternary epitope on trimeric BDBV GP, forms a high-affinity, slowly dissociating complex with BDBV GP and potently neutralizes pseudovirus carrying GP from both a 2007 reference strain and the currently circulating 2026 outbreak lineage. Importantly, in cynomolgus macaques, Maftivimab monotherapy initiated after the onset of clinical disease provided complete protection against lethal BDBV challenge, extending these *in vitro* findings to the gold-standard preclinical infection model. Authentic 2026-lineage virus is not yet available for challenge studies, but the conserved Maftivimab contact residues and similar neutralization of the 2007 and 2026 lineage support the expectation that the observed *in vivo* protection will hold against the 2026 lineage.

Using cryo-EM, we determined that Maftivimab engages a quaternary BDBV GP epitope which is equivalent to the epitope it engages on EBOV GP. Of note, residues 504–509, which vary among EBOV, BDBV-2007 and BDBV-2026, make at most weak contacts with Maftivimab in EBOV GP^11^ and are not resolved in our BDBV-2007 structure. This variation does not affect the activity of Maftivimab: it binds EBOV and BDBV-2007 GP with similar affinity and neutralizes pseudovirus carrying EBOV, BDBV-2007 or BDBV-2026 GP with similar potency. The N563 glycan also merits consideration: although residue N563 is not itself a direct Maftivimab contact, the N-glycosylation attached to it is directly contacted, and amino acid substitutions in N563 can confer resistance^11^. However, N563 is conserved between EBOV, BDBV-2007 and BDBV-2026, so this glycan contact is expected to be preserved in the current outbreak. Taken together, the conservation of the BDBV-2007 contact residues and the N563 glycan in BDBV-2026 GP indicates that the binding mode defined by our BDBV-2007 structure extends to the current outbreak lineage.

A recent computational study predicted that Maftivimab would be displaced from its canonical epitope on BDBV GP when modeled alone, but not when modeled within the full Inmazeb® cocktail^13^. Our experimental data show otherwise: a structure of Maftivimab bound alone to the BDBV GP trimer reveals engagement of the canonical epitope, consistent with its high-affinity binding, potent neutralization, and *in vivo* efficacy. This discrepancy likely reflects limitations of *in silico* models in capturing the multi-protomer geometry and glycan contacts of this epitope.

Antibody therapeutics developed against filovirus disease have typically combined virus neutralization with Fc-mediated effector functions^4,6,11,14^ — a paradigm shaped in part by early failures of purely neutralizing approaches: The potent human neutralizing antibody KZ52 failed to protect rhesus macaques despite high circulating antibody concentrations, suggesting that neutralization alone is insufficient for NHP efficacy against ebolaviruses^15^. This paradigm was challenged by BDBV289, a neutralizing glycan cap-directed mAb isolated from human survivors of the 2007 Uganda outbreak^16^. BDBV289 demonstrated that a purely neutralizing antibody can protect macaques against BDBV, though protection required two doses^17^. Here, we show that Maftivimab, a neutralizing-only antibody (previously shown to lack Fc effector activity)^6,11^ targeting the GP fusion loop region protected all treated cynomolgus macaques from lethal BDBV challenge with a single dose administered after the onset of clinical disease. To our knowledge, this is the first demonstration that a single-dose antibody acting exclusively through neutralization is sufficient to protect NHPs against an ebolavirus, establishing that potent neutralization of the viral fusion machinery alone is sufficient for NHP therapeutic efficacy, which may simplify future development of filovirus therapeutics.

Taken together, our data support the inclusion of Maftivimab in clinical trials as a monotherapy candidate for BDBV treatment during the ongoing outbreak and substantiate the WHO Technical Advisory Group’s May 2026 recommendation that it receive prioritized clinical evaluation against BDBV^18^. Beyond the current outbreak, our findings have implications for the future development of pan-ebolavirus or pan-filovirus antibody therapies.

## Methods

### Cryo-electron microscopy

#### REGN3479 IgG proteolysis

To facilitate cryo-electron microscopy structure determination, REGN3479 was cleaved into F(ab)’ fragments using IdeS FabRICATOR^19^. For simplicity of reading, “F(ab)’” will be referred to as “Fab” elsewhere in the manuscript. The cleavage reaction was performed at 37 °C for 30 min using 20 ng IdeS per 1 µg IgG and stopped by adding EDTA to a final concentration of 10 mM. Reduction was initiated by adding 6 mg 2-mercaptoethylamine (2-MEA) per 1 mL REGN3479 and incubating at 37 °C for 90 min. Reduced cysteines were capped by adding 10% of a 1 M iodoacetamide solution. Fc domains were removed using CaptureSelect IgG-Fc Affinity Matrix (Thermo Fisher Scientific), and the flow-through was further purified by fast protein liquid chromatography on a Superdex 200 Increase size-exclusion column (Cytiva). Fractions containing Fab fragments were pooled and concentrated.

#### Cryo-electron microscopy sample preparation

REGN3479 Fab was mixed in a 3:1 molar ratio with BDBV-2007 GP trimer and incubated on ice for 1 hour. Immediately prior to grid vitrification, n-Dodecyl-β-D-maltoside (DDM) was added to a final concentration of 0.13 mM (∼75% CMC). Quantifoil R 1.2/1.3 Cu 200 mesh grids were plasma cleaned in a Gatan 955 Solarus II, and 3 µL of the REGN3479:BDBV-2007 GP sample was added. After a 30 second incubation, the grids were blotted for 5 seconds with a blot force of 0 using a Vitrobot Mark IV (Thermo Fisher Scientific). Grids were stored in liquid nitrogen until data collection.

#### Cryo-electron microscopy data collection and processing

Electron micrographs were collected using Titan Krios transmission electron microscope (Thermo Fisher Scientific) operating at 300 kV and equipped with a K3 direct electron detector and BioQuantum energy filter (Gatan). 9,238 exposures were collected using EPU (Thermo Fisher Scientific) at pixel size of 0.839 Å and nominal defocus values from 0.5 to 2.2 µm. All exposures had a total flux of 47.3 e^-^/A^2^ and were fractionated into 40 frames. Data were processed using CryoSPARC v4.5.3^20^ (Structura). First, pre-processing was performed using patch motion correction and patch CTF estimation. Then, blob picker was used during the collection of the first ∼1,000 movies, along with concurrent 2D classification. Templates were selected from these 2D classes and used for template matching on all micrographs. A total of 334,505 particles were extracted, then curated using 2D classification. A final stack of 206,044 particles remained and were refined using non-uniform refinement^21^ to a gold-standard FSC resolution of 2.9 Å with imposed C3 symmetry.

#### Model building and refinement

An initial model of REGN3479 was obtained from the published structure of the Inmazeb cocktail bound to EBOV (PDB: 7TN9)^11^, and an initial model of the BDBV-2007 GP was predicted using Protenix v1.1.0^22^. The initial models were first rigid-body docked into the cryo-EM map using UCSF ChimeraX^23^ then iteratively refined in real space using Coot v0.9.8.95^24^ and PHENIX v2.2.1^25^. The structure was validated using the PHENIX implementation of MolProbity. All figures were generated using ChimeraX^23^.

### Antibody generation and surface plasmon resonance (SPR) binding kinetics

Maftivimab was generated using the VelocImmune® technology^6,9,10^. Binding kinetics between Maftivimab IgG and glycoprotein ectodomains from EBOV and BDBV were measured by surface plasmon resonance using a Biacore 8K instrument (Cytiva, Marlborough, MA) on a Series S CM5 sensor chip in filtered and degassed HBS-EP running buffer (10 mM HEPES, 150 mM NaCl, 3 mM EDTA, 0.05% (v/v) polysorbate 20, pH 7.4) at 37 °C. Both glycoprotein antigens were recombinantly expressed and purified in CHO cells with a C-terminal foldon trimerization domain and mmh (myc-myc-His) tag. The EBOV GP antigen comprised a mucin-like domain deletion construct spanning GP1 residues I33–V311 and GP2 residues T464–D637, with stabilizing substitutions T577P and K588F. The BDBV GP antigen comprised a mucin-like domain deletion construct spanning GP1 residues I33–P312 and GP2 residues I471–N643.

Capture sensor surfaces were prepared by covalently immobilizing a mouse anti-human Fc mAb (REGN2567) on the chip surface using the standard amine coupling chemistry, as reported previously^26,27^. Following surface activation, the remaining active carboxyl groups on the CM5 chip surface were blocked by injecting 1 M ethanolamine, pH 8.0 for 7 minutes. A typical resonance unit (RU) signal of ∼10,000 RU was achieved after the immobilization procedure.

Analysis of Maftivimab binding to glycoprotein ectodomains from EBOV (GenBank AHX24649.2) and BDBV-2007 (GenBank YP_003815435.1) was carried out by capturing the antibodies (∼300 RU) over immobilized anti-human Fc surface at 37 °C. Following the capture of the antibodies, different concentrations of glycoprotein ectodomains (25–100 nM, four-fold serial dilution) in the running buffer were injected for 1.5 minutes at a flow rate of 30 µL/min with a 5-minute dissociation phase. At the end of each cycle, the anti-human Fc surface was regenerated using a 12-second injection of 20 mM phosphoric acid.

All specific SPR binding sensorgrams were double-reference subtracted as reported previously^26^ and the kinetic parameters were obtained by globally fitting the double-reference subtracted data to a 1:1 binding model with mass transport limitation using Biacore™ Insight Evaluation software v6.0 (Cytiva, Marlborough, MA). The dissociation rate constant (kd) was determined by fitting the change in the binding response during the dissociation phase and the association rate constant (ka) was determined by globally fitting analyte binding at different concentrations. The equilibrium dissociation constant (KD) was calculated from the ratio of kd to ka. The dissociative half-life (t½) in minutes was calculated as ln2/(kd∗60).

### Epitope contact residue mapping

Contact residues of REGN3479 on BDBV-2007 GP were defined as residues with at least 15 Å² of buried surface area in the cryo-EM structure described above, and neighboring residues as those within 5 Å of a contact residue using UCSF ChimeraX^23^. GP amino-acid conservation was calculated from 60 BDBV glycoprotein sequences retrieved from NCBI GenBank^28^ (accessed 2026-10-01), deduplicated, retaining only near-complete genomes (>=18,000 nt, <=5% ambiguous bases, and no run of >10 consecutive N within the GP region). Sequences were aligned with MAFFT^29^ v7.526 against the BDBV reference genome NC_014373 and translated codon by codon; per-position conservation is reported as the percentage of sequences carrying the consensus residue shown in Figure 4.

### Pseudovirus neutralization assay

Neutralization potency of Maftivimab was evaluated using replication-incompetent vesicular stomatitis virus (VSV) pseudovirus encoding a luciferase reporter and carrying GP from EBOV (Makona, 2014, accession AHX24649.1), BDBV-2007 reference strain (YP_003815435.1), or BDBV-2026 (accession PP_006XCJJ.1; published May 17, 2026). An 11-point serial antibody dilution series was incubated with pseudovirus and applied to Vero cells for 24 h. Infection was quantified by luciferase activity, and % neutralization was calculated using uninfected control wells (cells and media; no pseudovirus, no mAb) and infected no-mAb control wells (cells, media and pseudovirus; no mAb), with the calculation 1-((well value-uninfected control)/(infected no-mAb control-uninfected control)). IC50, IC90, and IC99 values were derived by four-parameter logistic curve fitting using GraphPad Prism and are reported in Figure 1b.

### *In vivo* efficacy in cynomolgus macaques

Ethics statement: Animal research was conducted under an Institutional Animal Care and Use Committee (IACUC)-approved protocol (1972MF) in compliance with the Animal Welfare Act, Public Health Service (PHS) policy, and federal regulations related to experiments involving animals. Texas Biomedical Research Institute (Texas Biomed) is accredited by AAALAC International. All euthanasia was approved by a veterinarian and was performed using an overdose of sodium pentobarbital to minimize pain and distress in animals meeting predefined criteria for clinical assessment.

Ebolavirus H.sapiens-tc/UG/2007-200706291, CDC Seed virus #811250 (BDBV-2007 strain; NCBI accession MK028856.1) was isolated from a fatal human case in Uganda in 2007 and was obtained from Dr. Tom Ksiazek (University of Texas Medical Branch Health, Galveston National Laboratory) with a history of 2 cell culture passages in Vero E6 cells. BDBV-2007 was propagated with an additional cell culture passage at a multiplicity of infection of 0.001 in Vero E6 cells at Texas Biomedical Research Institute (TX Biomed) and used for NHP challenge experiments. Deep sequencing confirmed >99.99% sequence identity with MK028856.1 with a single partial polymorphism at position 17,198 that results in coding for both the reference leucine and phenylalanine at amino acid 1828 of the RNA-dependent RNA polymerase (L). Detection of a T -> TA frameshift at the 7U glycoprotein (GP) editing site occurred in approximately 7-8% of reads.

Healthy cynomolgus macaques (*Macaca fascicularis*) of Cambodian origin that were experimentally naïve were obtained from Inotiv, Inc. (West Lafayette, IN, USA) and were transferred to the Animal Biosafety Level 4 laboratory (ABSL4) at TX Biomed. Following acclimation, the animals were challenged intramuscularly (IM) with 1000 PFU of BDBV-2007. NHP weights ranged from 3.2 to 9.4 kg at the time of challenge and NHPs were 6.2 to 7.5 years of age. NHPs were randomized into treatment groups stratified by sex and by serum viral RNA load at 4 DPI. At 7 DPI, NHPs were treated by intravenous (IV) bolus of saline (placebo) or 150 mg/kg REGN3479 (n = 6 per group) at a dose volume of 2 mL/kg. Clinical observations occurred at least twice daily during ABSL4 acclimation through the end of the study, increasing to three or four times daily when signs of critical BDBV disease were observed. NHPs were followed for survival for 28 DPI; non-survival was defined as terminal illness based on cumulative clinical scoring as previously described^30^. Survival was compared between groups using a two-sided Boschloo’s exact test.

qRT-PCR was conducted to determine the viral load in serum. Briefly, serum was inactivated in TRIzol LS (Thermo Fisher Scientific) and RNA isolation was performed on a KingFisher Flex system (Thermo Fisher Scientific) with the NucleoMag Pathogen kit (Macherey-Nagel). RNA was quantified on a QuantStudio 5 Real-Time PCR System (Thermo Fisher Scientific) with TaqPath 1-Step RT-qPCR Master Mix (Applied Biosystems) with an initial step at 25°C for 2 minutes for primer annealing, 50°C for 15 minutes for reverse transcription, 95°C for 2 minutes for initial denaturation, and 45 cycles of 95°C for 3 seconds followed by 60°C for 30 seconds. qPCR probes (Integrated DNA Technologies) targeting BDBV GP were used: (Forward: 5’-TGA AGC CCC TGA GGG TGT AA-3’; Reverse: 5’-CCC TGT TCC AGA AAC CTT GTG-3’; Probe: 5’-FAM-AGG CTT CCC TCG CTG CCG TTA TG-TAMRA-3’). A plasmid containing the BDBV GP target sequence was synthesized (GenScript) and used to generate a standard curve used to calculate genomic equivalents (GE) per mL of serum, with a lower limit of detection of 2E+03 GE/mL.

## Acknowledgements

The work described in this manuscript was funded by Regeneron Pharmaceuticals and in part with federal funds from the U.S. Department of Health and Human Services (HHS); Administration for Strategic Preparedness and Response (ASPR); Biomedical Advanced Research and Development Authority (BARDA), under contract number HHSO100201700016C. The contract and federal funding are not an endorsement of the product or company. This investigation also used resources that were supported by the Southwest National Primate Research Center grant P51 OD011133 from the Office of Research Infrastructure Programs, National Institutes of Health. Research reported in this publication was supported by the Office of the Director, National Institutes of Health under Award Numbers S10OD028732 and S10OD032443. We thank the Texas Biomedical Research Institute Applied Science and Innovation Team for conducting *in vitro* and *in vivo* technical work.

